# Decidual innate immune cell kinetics following lipopolysaccharide challenge

**DOI:** 10.1101/2022.01.27.478092

**Authors:** Lauren E. St-Germain, Barbara Castellana, Jennet Baltayeva, Alexander G. Beristain

**Author notes:** To whom correspondence should be addressed: Alexander G. Beristain, British Columbia Children’s Hospital Research Institute, Department of Obstetrics & Gynecology, The University of British Columbia, Vancouver, British Columbia, Canada. V5Z 4H4.

## Abstract

In early pregnancy, macrophages (Mφ) and natural killer cells (NK) infiltrate and expand within the decidua to comprise 30% of all cellular content. These immune cell populations coordinate angiogenic and tissue remodeling processes that are needed for a healthy pregnancy. Importantly, decidual tissue-resident macrophages (trMφ) and uterine NK retain immunosurveillance properties that facilitate the targeting of infections (e.g., viral, bacterial). The timing and severity of these infections, as well as the resulting immune response, can dictate pregnancy outcome. However, little is known about the kinetics and activities of uterine myeloid and NK populations following infections. To address this knowledge gap, we defined the stepwise changes of uterine myeloid and NK subpopulations following lipopolysaccharide (LPS) challenge in a mouse model of early pregnancy. Low (25 µg/kg), moderate (50 µg/kg), and high (200 µg/kg) doses of LPS resulted in dose-dependent increases in peripheral and uterine inflammation, as well as a dose-dependent increase in the rate of fetal resorption. Compared with saline controls, mice exposed to LPS showed higher frequencies of immature monocytes, decreased TNFα-producing monocytes and Mφ, and increased conventional (c)NK expression of granzyme B in the uterus. These changes were followed by alterations in overall uterine (u)NK frequencies with increased cNK and decreased tissue resident (tr)NK. Together, this work describes how discrete levels of LPS-induced inflammation shape the innate immune cell landscape of the decidua. These findings establish insight into the stepwise immunological changes following endotoxin challenge and provide a better understanding of how inflammation controls the activity of key decidual leukocytes.

**Summary sentence:** Graded LPS challenge in early pregnancy leads to a stepwise increase in fetal resorption and associates with distinct alterations in frequencies and activities of uterine immune cells.

## INTRODUCTION

Optimal placental function and overall pregnancy success depend on tightly regulated angiogenic and vascular remodeling processes occurring within the decidual-fetal interface [1–3]. Early stages of uterine blood vessel growth, branching, pruning, and remodeling are controlled by diverse populations of leukocytes that infiltrate into and/or expand within the decidual microenvironment [4–9]. In addition to controlling vascular remodeling processes, uterine immune cells also function to protect the developing fetus and placenta from bacterial and viral infections. To this end, uterine immune cell activity must be appropriately regulated to enable placental development and uterine remodeling concurrent with the ability to mount timely and effective protection against infection.

Multiple subtypes of immune cells reside within the human decidual mucosa in early pregnancy, the most prevalent being monocyte-derived tissue-resident macrophages (trMφ), T cells, and a unique type of NK specific to the uterus, called uterine tissue-resident natural killer cells (trNK) [10–14]. Both trMφ and trNK are phenotypically and functionally different from their “conventional” immune cell counterparts found in circulating blood and other tissues. trMφ are heterogeneous; they are comprised of three distinct subtypes that can be defined by levels of CCR2 and CD11c expression and share features of M1- and M2-like Mφ states [15]. Similarly, three transcriptionally-defined subtypes of trNK are present in human decidual mucosa and all co-express the tissue-resident markers CD9 and CD49a [16]. In mice, decidual trMφ and trNK do not share the same complexity of surface markers or the diversity of defined cell subtypes as is known in humans; however, both decidual trMφ and trNK in humans and mice play conserved roles in spiral artery remodeling and uterine tissue angiogenesis, and generally promote a tolerogenic environment [17–21].

Cytokines, chemokines, and growth factors produced within the decidual-fetal interface influence leukocyte biology in the uterus. Aberrant inflammation, resulting from infection or pre- gestation health complications, can instruct decidual myeloid and NK populations to develop pro-inflammatory and pro-cytotoxic phenotypes that contribute to the development of, or exacerbate, severe pregnancy complications like spontaneous pregnancy loss, preterm birth, and preeclampsia [22–26]. [22–25]While aberrant inflammation associates with multiple complications of pregnancy, little is known about the specific cellular processes that are altered in the decidual-fetal interface due to inflammatory insult.

Modeling acute inflammation in mice via immune challenge with bacterial endotoxin lipopolysaccharide (LPS) in early pregnancy results in abrupt expression of multiple pro-inflammatory factors (e.g., TNFα and IL6) within utero-placental tissues [27]. This endotoxin-induced inflammation is known to associate with impaired uterine artery remodeling and fetal resorption; however, these effects are effectively blocked in IL15 knockout mice suggesting a central role for the IL15-dependent NK lineage in driving the effects of bacterial endotoxin exposure at the decidual-fetal interface [27]. Alternatively, while prolonged challenge to low and moderate doses of LPS in mid-late pregnancy in rats also results in impaired fetal growth and survival, these effects are essentially reversed in the presence of a TNFα antagonist [28–33]. This highlights the importance of mature monocytes and Mφ as central effectors following LPS challenge due to these cell types being dominant TNFα producers [34–36]. However, much remains to be understood about the decidual immune cell response following endotoxin exposure. These questions are further complicated by differences in immune cell composition that exist at different time-points of pregnancy in the dynamic decidual-fetal interface.

To examine the effects of endotoxin-instructed changes on uterine immune cell kinetics, pregnant mice were challenged with increasing doses of LPS to model a transient exposure to low, moderate, or high grades of inflammation. Exposure to all LPS doses led to robust induction of peripheral and local uterine inflammation. LPS dose positively correlated with rate of fetal resorption and LPS exposure induced transient changes in the frequencies of diverse types of myeloid cells as well as conventional and tissue-resident NK subsets in the uterus. Broadly, uterine immune cells responded to LPS challenge by increasing cellular granularity and cytokine expression. Immature monocytes and trMφ appeared first in the uterus in response to LPS exposure, followed by the activation and expansion of conventional natural killer cells (cNK). Taken together, this work describes how LPS modulates innate immune cell dynamics and activities within the decidual-fetal interface.

## METHODS

### Animals

Mouse experiments were carried out following the guidelines of the Canadian Council on Animal Care and approved by the UBC Animal Care Committee (protocol A18-0229). Female C57BL/6J and male DBA/2J mice were purchased from the Jackson Laboratory (Bar Harbor, ME, USA). Animals were housed under standard conditions in groups of maximum five mice in individually ventilated cages with water and food available ad libitum, under a 12:12 hrs light/dark cycle.

### Mouse model of LPS-induced inflammation in pregnancy

At 7 weeks of age, female C57BL/6J mice were paired with DBA/2J male studs. The appearance of a vaginal plug the next morning was taken as embryonic day (E) 0.5 of pregnancy. Because not all females become pregnant after first pairing, female age at conception ranged between 7-10 weeks. To induce a controlled and thoroughly defined type of inflammation in early pregnancy [27–36], pregnant female mice (dams) at E7.5 were injected intraperitoneally (i.p.) with LPS (*Salmonella typhimurium*; L6511, Sigma-Aldrich, St. Louis, MO, USA) in 100 µL saline at doses per unit of body weight reflecting low (10 µg/kg), moderate (25 µg/kg; 50 µg/kg) and high (200 µg/kg) doses of LPS challenge. As controls, dams were injected i.p. with 100 µL of saline. On E8.5, E9.5 and E17.5 females were euthanized, uterine/placental tissues were harvested, and blood was collected via cardiac puncture.

### Cytokine quantification

Whole blood was collected by saphenous vein blood draws 7 days pre-coitum at 7 weeks of age, 4 hrs post-injection (E7.5), and on E9.5 (via cardiac puncture). Serum was then purified from whole blood using serum separator tubes centrifuged for 5 min at 14000 rpm. Serum levels of TNFα, IL1β, IFNγ and IL6 were determined using a commercially available V-PLEX Mouse Cytokine 19-Plex assay (Meso Scale Discovery, Rockville, MD, USA) according to the manufacturer’s protocol.

### Whole blood WBC quantification

White blood cell (WBC) ratios (neutrophil/lymphocyte; monocyte/lymphocyte) were determined from whole blood collected by cardiac puncture at E9.5 using the Hemavet 950 FS multi-species hematology system (Drew Scientific, Germany).

### Gross fetal and placental tissue measurements

Implantation site intactness/resorption was determined at E9.5, where intact uterine horns were carefully dissected away from peritoneal connective tissue and imaged using a Nikon SMZ 745T triocular dissecting microscope (Nikon, Tokyo, Japan) outfitted with a digital camera. A dissection ruler was used for scale. At E17.5, implantation sites from E17.5 dams were dissected to remove the fetus and placenta/decidua, where tissues were imaged as above, and fetal length (mm) and weight (g), as well as placenta/decidua weight (g), diameter (mm), and area (mm^2^) were measured using ImageJ software (v1.53, NIH) or weighed (tissue wet weight) using an analytical scale. Fetal length was measured using crown-rump length, and placenta diameter was measured at its widest orientation.

### H&E, Immunohistochemistry (IHC), and immunofluorescence (IF) microscopy

Implantation sites from E9.5 and E17.5 were fixed in 4% paraformaldehyde for 24–48 hrs followed by processing into paraffin blocks. Histology, IHC, and IF workflow was performed as in Baltayeva et al. [37]. Serial tissue sections of approximately 6 µm were stained with hematoxylin and eosin (H&E) according to standard protocol. For IF staining tissues underwent antigen retrieval by heating slides in a microwave for 25 sec intervals for a total of 5 minutes at ∼90°C in a sodium citrate buffer (pH 6.0). Sections were then incubated in a blocking solution (5% bovine serum albumin in 1x phosphate-buffered saline (PBS) for 1 hr at room temperature (RT). Following blocking, samples were incubated with the following antibodies overnight at 4°C: rabbit polyclonal α-smooth muscle actin (SMA, 1:200; Abcam) and biotinylated DBA (1:100, Vector Laboratories). Slides were incubated with Alexa Fluor 568/488-conjugated streptavidin and goat anti-rabbit secondary antibody (1:200; Life Technologies) for 1h at RT. Glass coverslips were mounted onto slides using ProLong Gold Anti-Fade Reagent containing DAPI (Life Technologies). For each mouse, 2 implantation sites were evaluated (the 2^nd^ site from the midline on the anatomical right side of the uterine horn and 4th site from the midline on the anatomical left side of the uterine horn).

#### Artery remodeling measurements

Spiral artery remodeling measurements were performed on H&E-stained slides from dams sacrificed on E9.5 and E17.5. All spiral arteries within the decidual zone were measured in triplicate (three sections 48 µm apart from each other). Arterial wall size and lumen size was determined by measuring cross-sectional areas using ImageJ v1.53 software (NIH). Wall:lumen ratios were calculated by dividing the arterial wall area (µm^2^) over the arterial lumen area (µm^2^). Slides were imaged using 20× Plan-Apochromat/0.80 NA (E9.5) or 10x N-achroplan/0.25 NA (E17.5) objectives (Carl Zeiss, Jena, Germany) using an Axiocam 105 color digital camera (Carl Zeiss). SMA layer intactness was measured in spiral arteries of serially sectioned E9.5 implantation sites within the decidual zone. Following antibody labeling, slides were imaged using a 40× EC-Plan-Neofluar/0.9 PoI objective (Carl Zeiss). Arterial length and SMA layer were measured using ImageJ v1.53 software (NIH). SMA:perimeter ratios were calculated by dividing SMA+ vascular perimeter (µm) over arterial lumen perimeter (µm). IF images were obtained using an Axiocam 506 monochrome digital camera (Carl Zeiss). For each mouse, 2 implantation sites were evaluated with 3 sections obtained per implantation site at sections >48 µm apart from each other. At least 10 decidual arteries were assessed per section.

#### Implantation site microscopy imaging

H&E-stained slides of E17.5 implantation sites were used for placental morphology measurements. Total area (µm^2^) and areas of each individual placental layer (µm^2^) were measured using ImageJ v1.53 software (NIH). To normalize for differences in total placental areas, individual layer area was displayed as ratios of zone areas at each measurement point (i.e. labyrinth:junctional, labyrinth:decidual, and junctional:decidual ratios). Slides were imaged using 10× N-Achroplan/0.25 Ph1 objective (Carl Zeiss) using an Axiocam 105 color digital camera (Carl Zeiss).

### Decidual leukocyte preparations

For examination of uterine innate immune cell populations, murine implantation sites (∼7-8 implantation sites/mouse) were first separated from uterine tissues before being mechanically and enzymatically processed to obtain single-cell suspensions. In brief, tissues were washed in cold 1X PBS (pH 7.4). Following this, tissues were minced with a razor blade and subjected to a 30-minute enzymatic digestion at 37 °C in 3 mL of 1:1 DMEM/F12 media (Gibco, Grand Island, NY; 200 mM L-glutamine) with 1X-collagenase/hyaluronidase (10X stock; StemCell Technologies, Vancouver, Canada), 80 μg/mL DNaseI (Sigma, St. Louis, MO), 1% penicillin/streptomycin, and 1X antibiotic–antimycotic solution (100X dilution, Gibco). Single-cell suspensions were passed through a 100 µm strainer and decidual leukocytes were enriched using a Percoll (GE Healthcare) density gradient. Decidual leukocytes were immediately used for cell-surface marker characterization via flow cytometry analysis.

### Flow cytometry

For flow cytometry, 1 x 10^6^ isolated leukocytes were either collected immediately after single cell isolation or cultured for a total of 5 hrs in complete medium (RPMI1640 medium complemented with 10% FBS, 1% Penicillin/Streptomycin, 1 mM Sodium Pyruvate, and 55 nM of 2-Mercaptoethanol) with the protein transport inhibitor Brefeldin A (Thermo Fisher) added during the last 4 hrs of incubation at 37°C and 5% CO_2_. Supernatants were collected with remaining adherent cells collected by 5 min incubation with Cell Dissociation Buffer (Gibco, Thermo Fisher Scientific) at 37°C and 5% CO_2_. Cells were then collected and centrifuged at 1,200 rpm for 5 min at 4°C and pellets were washed in 1X PBS, followed by incubation with Fix Viability dye 780 (1:1000; Thermo Fisher Scientific) for 15 min at RT, protected from light. After, cells were washed and pelleted down by centrifugation and resuspended in anti-mouse CD16/CD32 (eBiosciences, Thermo Fisher Scientific) diluted to 1:25 in Super Brilliant Stain Buffer (BD Biosciences) for 10 min at 4°C. Next, cells were stained for the following extracellular markers: anti-CD3ε APC ef780 (1:50; clone 145-2C11), anti-CD19 APC ef780 (1:50; clone 1D3), anti-CD45 ef450 (1:150; clone 30-F11), anti-CD49b PeCy7 (1:100; clone DX5), anti-NK1.1 FITC (1:50; clone PK136), anti-F4/80 BV780 (1:100; clone BM8) from eBiosciences, anti-CD11c PeCF594 (1:100; clone N418; BD Biosciences), and anti-CD49a APC (1:50; clone HMα1) and anti-CD11b BV570 (1:100; clone M1/70) from BioLegend. Cells were fixed for 30 min at 4°C using eBioscicences^TM^ FoxP3/Transcription factor staining buffer set (Thermo Fisher) and stained for the following intracellular markers: anti-IFN*γ* AlexaFluor 700 (1:50; clone XMG1.2; eBiosciences), anti-granzyme B PE (1:50; clone QA16A02) and anti-TNFα BV711 (1:50; clone MP6-XT22) from BioLegend in permeabilization buffer at 4°C for 30 min. After staining, cells were washed and resuspended in 200 uL of 1X PBS with 2% FBS. Data was acquired using a BD LSRFortessa X-20 (BD Biosciences) and analysed by FlowJo 10.7.2 software (Tree Star, Inc., Ashland, OR, USA).

### RNA extraction

Prior to RNA extraction, the 2^nd^ uterine implantation site from midline on the anatomical right side was snap frozen in liquid nitrogen. The snap-frozen tissue was then biopulverized under super-cooled conditions, and total RNA was isolated using TRIzol reagent (Life Technologies) according to the manufacture’s protocol. RNA was further purified using RNeasy MinElute Cleanup columns (Qiagen), and RNA purity and concentration (ng/μl) was confirmed using a NanoDrop Spectrophotometer (Thermo Fisher Scientific) where a 260/280 readout greater than or equal to 1.8 was considered pure.

### cDNA synthesis and qPCR

RNA was reverse-transcribed using a first-strand cDNA synthesis kit (Quanta Biosciences) and subjected to qPCR (ΔΔCT) analysis using PowerUp™ SYBR™ Green Master Mix (Thermo Fisher Scientific) on an ABI ViiA 7 Real-Time PCR system (Thermo Fisher Scientific). Forward and reverse primer sets were designed using NCBI Primer-Blast software and purchased from Integrated DNA Technologies. Forward and reverse primer sets are as described: *Il6* (F: 5’- GCTGGAGTCACAGAAGGAGTGGC -3’, R: 5’- GGCATAACGCACTAGGTTTGCCG -3’); *Il1b* (F: 5’- TGGACCTTCCAGGATGAGGACA-3’, R: 5’- GTTCATCTCGGAGCCTGTAGTG-3’); *Tnf* (F: 5’- GTAGCCCACGTCGTAGCAAA -3’, R: 5’- ACAAGGTACAACCCATCGGC -3’); *Ifng* (F: 5’- AGACAATCAGGCCATCAGCAA-3’, R: 5’- GTGGGTTGTTGACCTCAAACT-3’); *Gzmb* (F: 5’- ACCCAAAGACCAAACGTGCT -3’, R: 5’- TTTTCCCCTCCCACTTAGCC -3’); *Gzmb* (F: 5’- AGCTACTGAAGCAAGGCACA -3’, R: 5’- AAGCCAATGCTCTGACAAGGT -3’); *Prf1* (F: 5’- GCATGGGAGAAAGGAACTGGA -3’, R: 5’- ACTTCACGCCACTATTCCAGAG -3’); *Hprt* (F: 5’- CTGGTGAAAAGGACCTCTCG-3’, R: 5’-TGAAGTACTCATTATAGTCAAGGGCA-3’) [38]. All raw data was analyzed using StepOne Software (V2.3; Applied Biosystems: StepOnePlus). Threshold cycle (Ct) values were used to calculate relative gene expression levels. All values were normalized to *HPRT* transcripts.

### Statistical analysis

Data was assessed using either one-way or two-way ANOVA with Tukey’s multiple comparison test and single pooled variance. *P* <0.05 was taken as the threshold for significance. All statistical analyses were performed using Prism9 software (GraphPad Software, Inc.).

## RESULTS

### LPS challenge in early pregnancy transiently drives peripheral and uterine-specific inflammation

To understand how decidual innate immune cell populations are affected by inflammatory insult, we established an acute model of bacterial endotoxin exposure in early pregnancy. To do this, three distinct concentrations of LPS (*Salmonella typhimurium*) representing low- (25 μg/kg), moderate- (50 μg/kg), and high- (200 μg/kg) grades of inflammatory challenge were injected i.p. on embryonic day (E)7.5 in C57BL/6J dams (Figure 1A). We chose this gestational timepoint because at this time the placenta is still developing, is non-functional, and is highly susceptible to inflammatory insults [37, 39]. Further, decidual immune cell content is expanding substantially at this stage [13] indicating that LPS challenge will likely have its greatest effects on major immune cell populations found in the decidua during this developmental window. Lastly, C57BL/6J dams were paired with DBA/2J males to better reflect genetic heterogeneity of the conceptus seen in human pregnancies. Three temporal assessments of both peripheral and local tissue inflammation were assessed (at 7 days pre-coitum, on E7.5, and on E9.5; Figure 1A).

**Figure 1.**
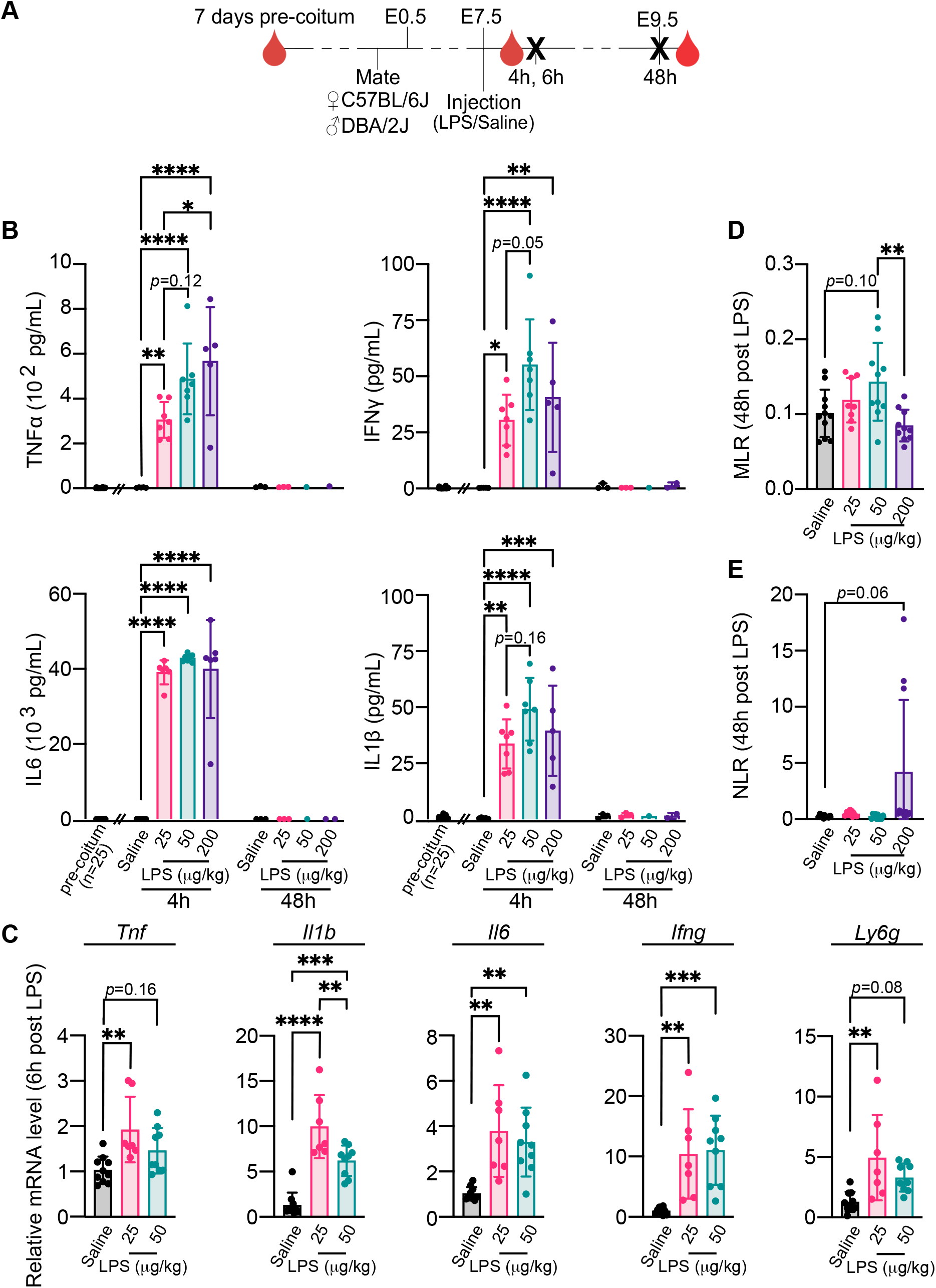
Single LPS challenge in early pregnancy induces transient peripheral and uterine inflammation. (A) Schematic of timeline for mouse mating, injection, blood collection (red blood droplet), and sacrifice (bold **X**) for Figure 1 experiments. (B) Serum concentrations of inflammatory factors TNFα, IFNγ, IL6, and IL1β; 7 days pre-coitum (n = 25), 4 hrs after injection (4 hrs post-inject; Saline, *n* = 5; 25 µg/kg LPS, *n* = 7; 50 µg/kg LPS, *n* = 7; 200 µg/kg LPS, *n* = 6), and 48 hrs after injection (48 hrs post-inject; Saline, *n* = 3; 25 µg/kg LPS, *n* = 3; 50 µg/kg LPS, *n* = 1; 200 µg/kg LPS, *n* = 2). (C) Gene expression levels of *Tnf*, *Il1b*, *Il6*, *Ifng*, and *Ly6g* within uterine tissue 6 hrs after injection. Saline, *n* = 8; 25 µg/kg LPS, *n* = 6; 50 µg/kg LPS, *n* = 7. (D) Monocyte: lymphocyte (MLR) and (E) neutrophil: lymphocyte (NLR) ratios from whole blood collected 48 hrs following injection, measured using the Hemavet 950 FS multi-species hematology system. Saline, *n* = 11; 25 µg/kg LPS, *n* = 9; 50 µg/kg LPS, *n* = 10; 200 µg/kg LPS, *n* = 11 mice. Data is represented as mean ± SD. Differences between groups were compared using one-way (C-E) and two-way (B) ANOVA. **\*** ≤ 0.05, ** ≤ 0.01, *** ≤ 0.001, **** ≤ 0.0001.

To measure the level of peripheral inflammation induced by all three doses of LPS-challenge, we quantified the expression of 4 pro-inflammatory cytokines [40–44] in maternal blood 4 hrs after LPS exposure using an enzyme-linked immunosorbent assay. Compared to control mice injected with saline, injections of low, moderate, and high concentrations of LPS led to significant increases in serum TNFα, IFNγ, IL6, and IL1β 4 hrs following challenge; TNFα and IFNγ showed LPS dose-dependent increases (Figure 1B). In control mice, baseline levels of these cytokines remained constant and markedly lower than in LPS challenged mice; E7.5 cytokine levels did not differ between pre-coitum levels and E9.5 levels in saline-injected mice (Figure 1B). Examining corresponding mRNA transcripts in uterine tissue 6 hrs post LPS challenge by qPCR showed similar readouts to what we observed systemically. Though tissue *Tnf* levels increased following challenge with only low but not moderate LPS dose, mRNA levels of *Ifng, Il6,* and *Il1b* were on average 4-fold greater in both low and moderate LPS treatments compared to controls (Figure 1C).

We also assessed levels of *Ly6g* transcripts, encoding for Lymphocyte antigen 6 complex locus G6D (Ly6G), which are predominately expressed by neutrophils [45], as a readout of neutrophil- content and inflammation. Similar to *Tnf,* low-dose LPS led to increased levels of *Ly6g* mRNA levels in uterine tissue, suggesting that pro-inflammatory neutrophils are present in the decidua of LPS- challenged mice. However, the moderate LPS dose resulted in a modest, but not significant increase in *Ly6g* when compared to Saline-injected mice (Figure 1C). As a positive control for neutrophil infiltration and *Ly6g* expression, we separately examined *Ly6g* transcripts in intestinal tissue samples collected from an inflammatory bowel disease (IBD) mouse model shown to have abundant neutrophil infiltration in the intestine [46]; as expected *Ly6g* levels were > 3000-fold greater than those observed in uterine tissue from control mice injected saline (Supplemental Figure 1A). Importantly, transcript levels (of *Tnf*, *Ifng*, *Il6*, *Il1b*, and *Ly6g*) were not measurable in uterine tissue collected from dams that were exposed to high dose LPS because of high tissue resorption (> 80%) and inconsistent uterine morphology that impaired tissue identification and extraction (see below).

Cellular readouts of peripheral inflammation were also examined 48 hrs post LPS injection by measuring whole blood monocyte/lymphocyte (MLR) and neutrophil/lymphocyte (NLR) ratios in a sub-cohort of mice. Consistent with previous reports that describe subclinical types of inflammation [47]utrophil/lymphocyte (NLR) ratios in a sub-cohort of mice. Consistent with previous reports that describe subclinical types of inflammation [47], moderate LPS challenge showed a moderate, but non- significant, increase in the MLR compared to control mice (Figure 1D). While the M[48–51] (Figure 1E). Together, these data indicate that dose-dependent one-time i.p. injections of LPS at E7.5 recapitulate a defined type of inflammation showing incremental increases in specific inflammatory readouts.

### LPS exposure drives fetal resorption in a dose-dependent and bimodal manner

Having characterized systemic and uterine tissue inflammatory readouts, we next examined how stepwise increases in LPS affect fetal survival and growth. LPS challenge resulted in increased rates of fetal resorption; notably, this response was bimodal and dose-dependent (Figure 2B, 2C, 2D). At E9.5 in saline-injected control dams, 2 of 9 mice showed resorption rates between 10% and 15% of implanted conceptuses; the remaining 7 control dams did not show indications of fetal resorption (Figure 2B, 2C). In contrast to this, LPS challenge resulted in increased rates of fetal resorption; notably, this response was binary and dose-dependent (Figure 2B, 2C). Specifically, LPS exposure led to either low/no rates of resorption that mirrored the rates observed in saline controls, or showed complete (i.e., 100%) resorption of all implantation sites (Figure 2C). This binary response skewed more towards complete resorption as LPS dose increased, where resorption rates of 22% (2/9), 58% (7/12), and 82% (9/11) were observed in low-, moderate-, and high-dose LPS conditions, respectively (Figure 2C).

**Figure 2.**
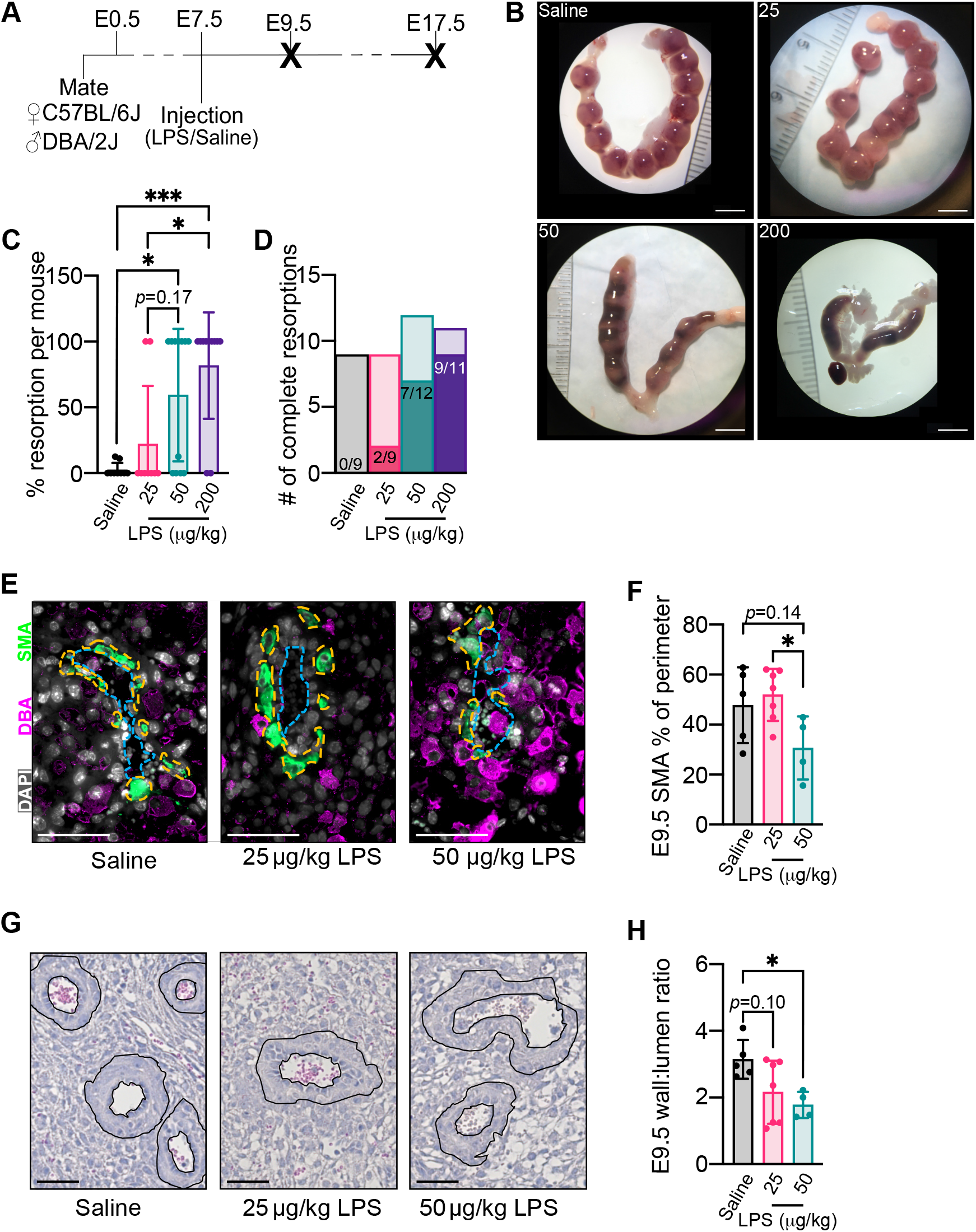
Stepwise increase in fetal resorption observed after graded LPS injection as well as transient enhancement of arterial remodeling. (A) Schematic of timeline for mouse mating, injection, and sacrifice (bold X) for Figure 2 experiments. (B) Representative images of uterine horns 48 hrs following injection. Scale bar = 5 mm. (C) Percentage of resorbed implantation sites per mouse 48 hrs following injection. (D) Numbers of pregnancies with complete resorption n E9.5 for each LPS dose. Saline, *n* = 6; 25 µg/kg LPS, *n* = 5; 50 µg/kg LPS, *n* = 4 mice. (E) Representative images of E9.5 arteries stained for DAPI, SMA, and DBA+. Scale bars = 50 µm. Yellow dashed lines describe the measurement of SMA layer. (F) Proportion of smooth muscle actin in artery perimeter per mouse at E9.5. Saline, *n* = 5; 25 µg/kg LPS, *n* = 7; 50 µg/kg LPS, *n* = 4 mice. (G) Representative images of arteries from E9.5 implantation sites 48 hrs following injection, stained for hematoxylin and eosin. Scale bars = 50 µm. (H) Wall:lumen ratio per mouse at E9.5, 48 hrs following injection. Saline, *n* = 5; 25 µg/kg LPS, *n* = 7; 50 µg/kg LPS, *n* = 4 mice. Data is represented as mean ± SD. Differences between groups were compared using one-way ANOVA. * ≤ 0.05, ** ≤ 0.01, *** ≤ 0.001, **** ≤ 0.0001.

While LPS challenge increased fetal death at E9.5, LPS challenge in the surviving fetuses did not affect implantation site number or size compared to controls (Supplemental Figure 2B, 2C). Similarly, in surviving pregnancies at E17.5, LPS treatment did not impact fetal or placental weight, nor did it affect the fetal/placental weight ratio (Supplemental Figure 2D, 2E, 2F). Likewise, fetal crown-rump length and placental diameter were also not affected by either low- or moderate-dose LPS (Supplemental Figure 2G, 2H). Examining placental and decidual zones at E17.5 in both control and LPS-challenged mice overall showed that LPS exposure did not affect placental development (Supplemental Figure 3A, 3B, 3C, 3D). However, exposure to low amounts of LPS did lead to slightly larger labyrinth areas than in control and moderate LPS dose mice, and this was reflected by moderately larger placental areas, though these effects were not significant (Supplemental Figure 3B, 3C). Together, these findings show that dose-dependent exposure to LPS impairs fetal/implantation site survival through a bimodal response. Notably, however, in pregnancies that survive even high-dose LPS challenge, gross fetal and placental metrics appear normal.

### LPS-induced inflammation associates with enhanced uterine artery remodeling

In previous studies, exposure to LPS affects utero-placental vascular remodeling processes important in pregnancy. Because a substantial proportion of low- and moderate LPS-challenged mice did not result in resorption and showed minimal histological alterations of the placenta, we next set out to measure the effect of LPS on uterine artery remodeling. To measure artery remodeling, E9.5 implantation sites from saline- and LPS-challenged dams were probed with antibody targeting SMA. In saline-injected dams, the SMA perimeter was approximately 50% of uterine artery area, suggesting that artery remodeling was taking place (Figure 2F). The observed SMA intactness in mice exposed to low-dose LPS challenge was not visibly different from that observed in mice injected with saline (Figure 2F). In contrast, moderate dose LPS treated mice showed an enhancement of fragmented SMA signal than in controls and a reduction in SMA perimeter around vessels, though this difference was not statistically significant (Figure 2F). However, comparing SMA intactness between dams exposed to low and moderate dose LPS challenge showed that remodeling processes were greater in mice exposed to moderate-dose LPS challenge versus mice exposed to low-dose LPS challenge (Figure 2E, 2F).

Vascular remodeling was further assessed by histology using H&E staining on E9.5 implantation site deciduae to measure artery wall thickness (Figure 2G). Compared to saline injected control dams, both low and moderate concentrations of LPS decreased uterine artery wall:lumen ratios, though the low dose treatment did not meet statistical significance (Figure 2H). To assess if these physiological differences in arterial remodeling driven by LPS are maintained in late pregnancy, E17.5 implantation site artery wall thicknesses were examined. In saline control dams, wall:lumen ratio was decreased compared to the E9.5 time-point (Figure 2H). However, at this later developmental period, LPS challenge did not further potentiate artery wall remodeling, suggesting that LPS imparts only a transient acceleration in vasculature remodeling close to the time of challenge (Supplemental Figure 4A, 4B).

### LPS differentially impacts decidual myeloid and NK composition

Because decidual myeloid and NK subsets play central roles in orchestrating uterine tissue and vascular remodeling, we next set out to examine if LPS exposure affects these key innate immune populations of the uterus. To examine the effects of LPS challenge in modulating major populations of decidual myeloid and NK immune cells, we focused our analyses on low- and moderate-dose LPS challenge, as high-dose LPS rarely produced viable implantation sites needed to accurately isolate. To assess immune cell kinetics following LPS challenge, we immunophenotyped diverse subsets of decidual myeloid [immature monocytes (Mo); mature Mo/Mφ; trMφ; and myeloid-derived dendritic cells (mDC)] and uterine (u)NK [tissue resident NK (trNK); conventional NK (cNK)] subtypes at 6 and 24 hrs post-LPS challenge on E7.5 (Figure 3A). For multicolor flow cytometry phenotyping, broad decidual myeloid and uNK subsets were identified by negative lineage gating of CD3ε (T cells) and CD19 (B cells) and through positive gating of CD11b (myeloid) and NK1.1 (NK) lineage markers (Figure 3B). While total myeloid and uNK cell frequencies were not affected by LPS exposure, overall myeloid frequencies naturally increased from ∼30% to ∼60% between E7.5 and E8.5, respectively (Figure 3C).

**Figure 3.**
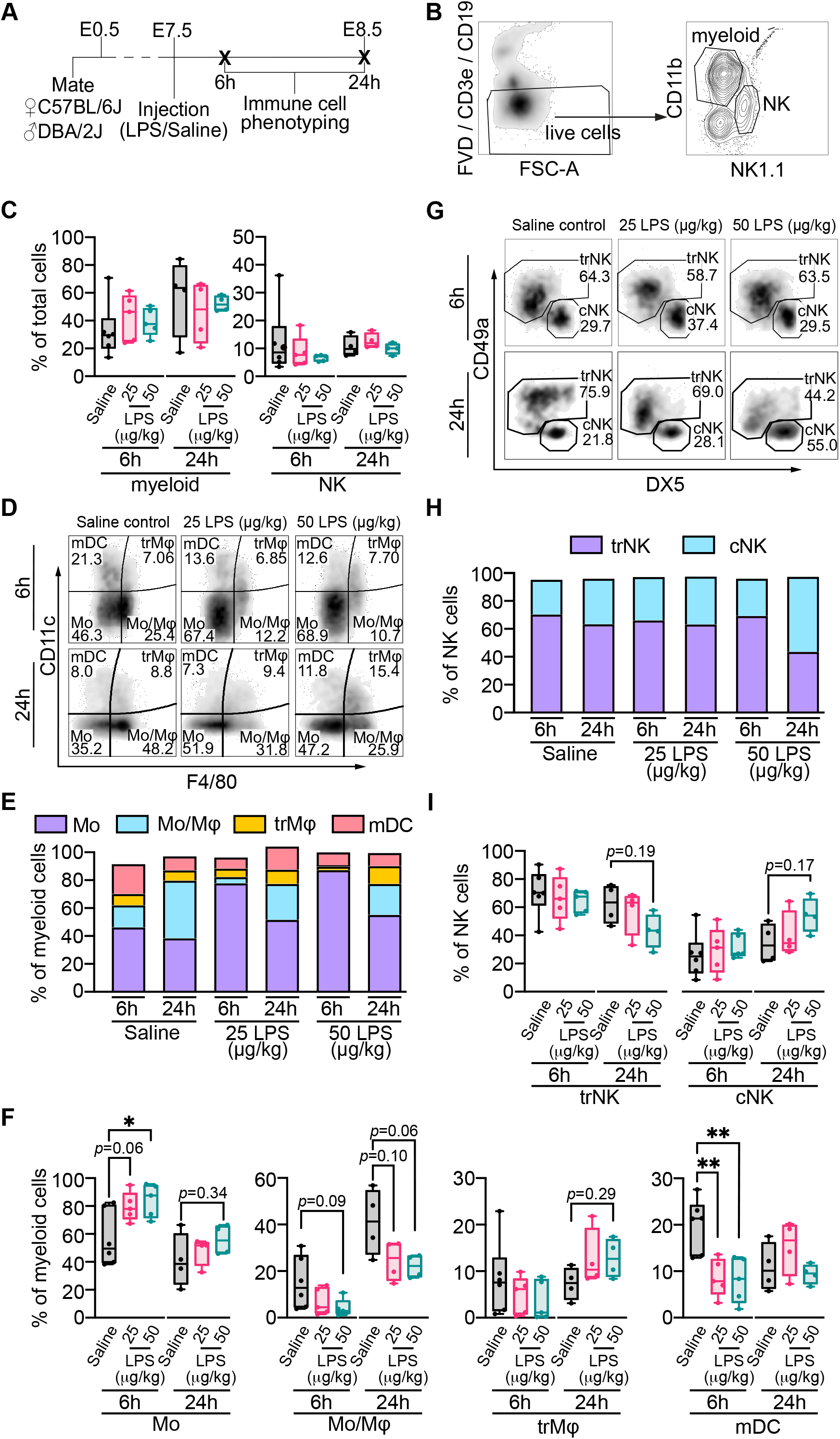
Abrupt uterine innate immune response to LPS challenge. (A) Schematic of timeline for mouse mating, injection, and sacrifice (bold **X**) for immunophenotyping experiments (Figures 3-6). (B) Gating strategy used to identify myeloid and NK populations from live, CD3ε^-^CD19^-^ cells. (C) Myeloid and NK as a percentage of total live cells. (D) Representative gating of myeloid populations: mDC, trMφ, immature monocytes (Mo), and mature monocytes/Mφ (Mo/Mφ) for different LPS doses (Saline, 25 µg/kg, 50 µg/kg), and 2 time points. (E) Median myeloid populations as proportions of total myeloid cells. (F) Myeloid populations as proportions of total myeloid cells. (G) Representative gating of NK populations (trNK, cNK) for different LPS doses, and 2 time points. (H) Median NK populations as proportions of total NK. (I) NK populations as proportions of total NK. The 6 hrs and 24 hrs timepoints are measured using two separate cohorts of mice. For the 6 hrs timepoint: Saline, *n* = 8; 25 µg/kg LPS, *n* = 6; 50 µg/kg LPS, *n* = 7 mice. For the 24 hrs timepoint: Saline, *n* = 4; 25 µg/kg LPS, *n* = 4; 50 µg/kg LPS, *n* = 4 mice. Data is represented as box plots with median ± min/max and interquartile range (C, F, I) or as summary data showing only the median values (E, H). Differences between groups were compared using two-way ANOVA. **\*** ≤ 0.05, ** ≤ 0.01, *** ≤ 0.001, **** ≤ 0.0001.

We examined specific myeloid subsets using the CD11c and F4/80 markers to distinguish between immature monocytes (Mo) (F4/80^-/^CD11c^-^), mature Mo/Mφ (F4/80^+^/CD11c^-^), trMφ (F4/80^+^/CD11c^+^), and mDC (F4/80^-^/CD11c^+^) (Figure 3D). Six hrs post LPS challenge, both the low- and moderate-dose LPS challenges led to substantial increases in the frequency of immature monocytes (Figure 3E, 3F). This increase in the proportion of monocytes at 6 hrs post LPS exposure was reciprocally characterized by decreases in frequencies of mature Mo/Mφ and mDC (Figure 3E, 3F). Notably, LPS dose did not potentiate differences in myeloid sub-populations, where both low and moderate LPS treatment imparted similar effects. At E8.5, 24 hrs post LPS exposure, similar frequencies of Mo and DCs were observed in saline control and LPS groups (Figure 3F). This contrasted with the sustained decrease in mature decidual Mo/Mφ frequency that was observed in LPS-challenged mice (Figure 3F). Moreover, a moderate increase in the proportion of trMφ was seen at 24 hrs post-LPS challenge, though these differences were not statistically significant (Figure 3F).

Focusing on trNK (CD49a+) and cNK (DX5+) subpopulations of uNK (Figure 3G), LPS challenge did not affect the frequencies of either subset 6 hrs post-challenge (Figure 3H, 3I). However, 24 hrs post LPS challenge (E8.5), the moderate LPS dose did result in an approximate 2-fold increase in frequency of cNK, though these differences were not significant (Figure 3H, 3I). The increase in proportion of cNK at 24 hrs in mice receiving moderate LPS dose was also reflected in an overall decrease in the frequency of trNK; in control mice, trNK comprised >60% of uNK, whereas in mice injected with moderate LPS dose the proportion of trNK fell to 40% (Figure 3I). Alterations in cNK and trNK frequencies were not as evident in low-dose challenge, though a trend for decreased trNK frequency at 24 hrs post treatment was seen (Figure 3I). Together, these data show that LPS initially drives alterations in myeloid lineage cell populations as early as 6 hrs post challenge. Following these early cellular changes, alterations in cNK and trNK subsets are observed 24 hrs later that may reflect either the expansion of uterine cNK or the infiltration of peripheral blood NK.

### LPS engages distinct states of innate immune cell activity

Because proportions of decidual myeloid and NK populations were altered following LPS challenge, we next set out to examine how LPS might also affect the timing of functional readouts of cytokine production and cytotoxicity. To do this, LPS challenge was induced at E7.5 as before, and immune cells from implantation sites were harvested at either 6 or 24 hrs post LPS treatment. Cells were then immunophenotyped for the intracellular protease granzyme B, a factor important in orchestrating apoptosis [52], and the cytokines TNFα and IFNγ, important in inflammatory responses in infection and in mediating uterine tissue and vascular remodeling processes important in pregnancy [53–58] (Figure 4A). For these analyses, trMφ were not assessed due to inconsistences in capturing adequately powered flow cytometry events.

**Figure 4.**
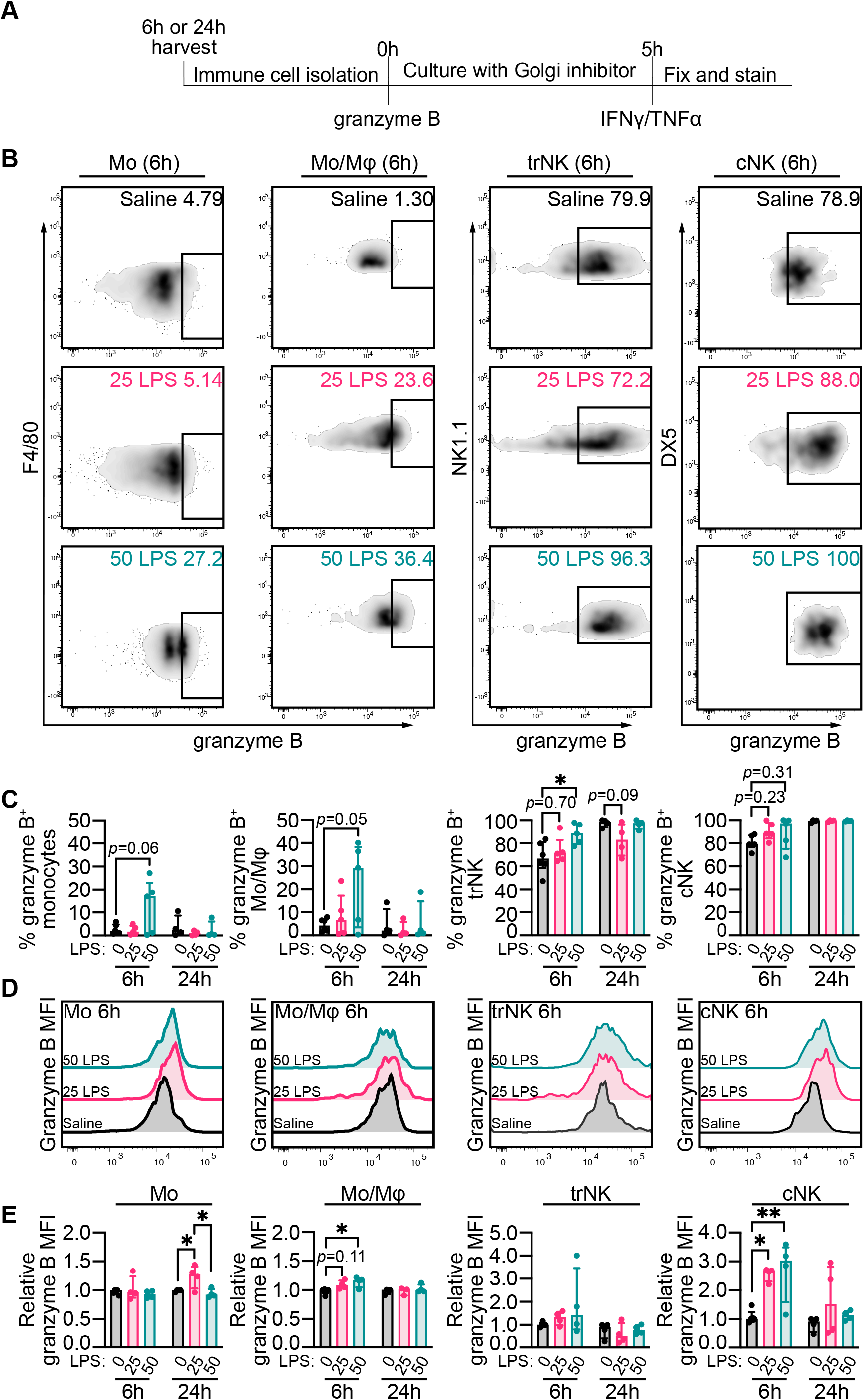
Temporal kinetics of uterine myeloid and NK intracellular cytolytic factor levels following LPS challenge. (A) Schematic of timeline for uterine tissue processing, starting with the harvest of uterine horns from dams 6 hrs and 24 hrs following injection with LPS or Saline control. (B) Representative gating for expression of intracellular granzyme B 6 hrs following exposure to LPS (Saline, 25 µg/kg, 50 µg/kg) in immature monocytes (Mo), mature monocytes/Mφ (Mo/Mφ), trNK, and cNK in the decidua. (C) Proportions of granzyme B^+^ Mo, Mo/Mφ, trNK, and cNK 6 hrs and 24 hrs post-injection. (C) Mo, Mo/Mφ, trNK, and cNK expression of granzyme B 6 hrs post-injection. (D) Median fluorescent intensity (MFI) of granzyme B in Mo, Mo/Mφ, trNK, and cNK (values relative to a Saline control for each experimental round) 6 hrs and 24 hrs post-injection. The 6 hrs and 24 hrs timepoints are measured using two separate cohorts of mice. For the 6 hrs timepoint: Saline, *n* = 8; 25 µg/kg LPS, *n* = 6; 50 µg/kg LPS, *n* = 7 mice. For the 24 hrs timepoint: Saline, *n* = 4; 25 µg/kg LPS, *n* = 4; 50 µg/kg LPS, *n* = 4 mice. Data is represented as mean ± SD. Differences between groups were compared using two-way ANOVA. **\*** ≤ 0.05, ** ≤ 0.01, *** ≤ 0.001, **** ≤ 0.0001.

Flow cytometric assessment of intracellular granzyme B showed that moderate concentrations of LPS resulted in increased frequencies of granzyme B+ in immature Mo and mature Mo/Mφ 6 hrs post challenge, though these increases were not significant (Figure 4B, 4C). On the other hand, while proportions of granzyme B^+^ trNK approach 70% in control mice, proportions of granzyme B^+^ trNK approached 90% in mice exposed to moderate-LPS challenge (Figure 4B, 4C). In saline controls, 80% of cNK expressed granzyme B, and low and moderate LPS treatment increased the frequency of cNK expressing granzyme B to ∼100% (Figure 4B, 4C). Intracellular levels of granzyme B, as measured by median fluorescence intensity (MFI), were increased 6 hrs post treatment of moderate grade LPS in immature monocytes and activated Mφ (Figure 4D, 4E). However, this increase was no longer seen 24 hrs post treatment (Figure 4D, 4E). Similar to proportions of trNK expressing granzyme B, MFI levels of granzyme B following LPS engagement were not altered (Figure 4D, 4E). This contrasted with cNK that showed strong induction of granzyme B 6 hrs following low and moderate LPS challenge; by 24 hrs post challenge, granzyme B levels were indistinguishable between saline controls and LPS groups (Figure 4D, 4E).

To assess intracellular levels of TNFα and IFNγ in decidual monocytes and Mφ, uterine single cell isolates were cultured for 5 hrs in brefeldin A (a potent protein trafficking inhibitor) and stained for flow cytometric readouts (Figure 5A). LPS treatment led to overall reductions in the proportions of TNFα-expressing Mo/Mφ from 70% (saline) to 40% (LPS; low and mid dose) 6 hrs post challenge (Figure 5A, 5B). Consistent with these changes, MFI levels of TNFα in both immature Mo (24 hrs) and mature Mo/Mφ (6 and 24 hrs) were decreased following LPS challenge (Figure 5C, 5D). In contrast, trNK (6 hrs and 24 hrs post-challenge) and cNK (24 hrs post-challenge) showed moderate increases in TNFα^+^ frequencies following LPS exposure, though these differences were not significant (Figure 5E, 5F). Changes in frequencies of either uNK subtype expressing TNFα were not reflected by increases in MFI (Figure 5G, 5H).

**Figure 5.**
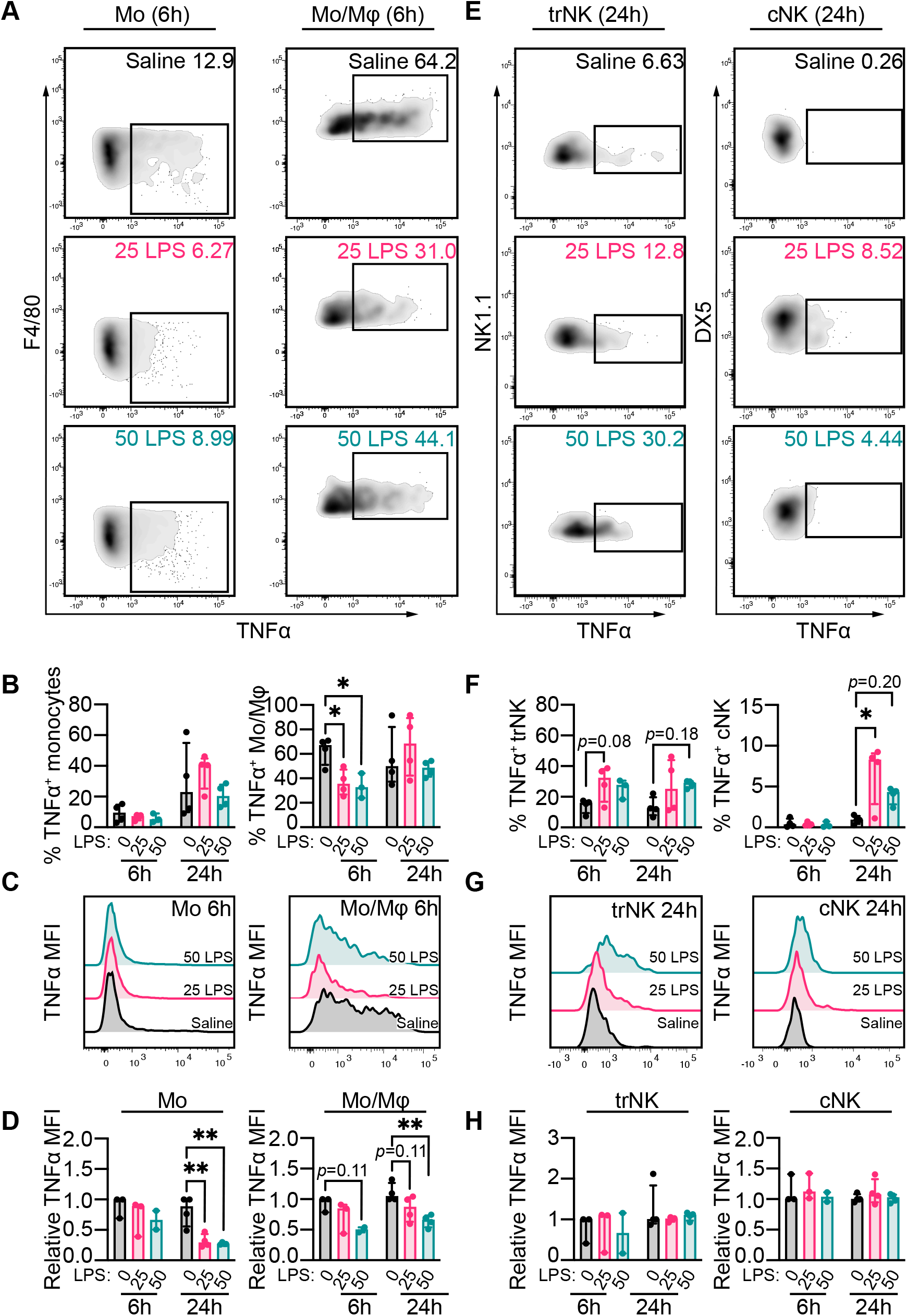
Temporal kinetics of uterine myeloid and NK intracellular TNFα levels following LPS challenge. (A) Representative gating for expression of intracellular TNFα 6 hrs following exposure to LPS (Saline, 25 µg/kg, 50 µg/kg) in immature monocytes (Mo), mature monocytes/Mφ (Mo/Mφ), trNK, and cNK in the decidua. (B) Proportions of IFNγ^+^ and TNFα^+^ Mo and Mo/Mφ 6 hrs and 24 hrs post-injection. (C) Mo and Mo/Mφ expression of IFNγ and TNFα, 6 hrs post-injection. Biological control for Mo expression of IFNγ and TNFα is non-cultured monocytes, biological control for Mo/Mφ expression of IFNγ and TNFα is non-cultured Mo/Mφ. (D) Median fluorescent intensity (MFI) of IFNγ and TNFα in Mo and Mo/Mφ (values relative to a Saline control for each experimental round), 6 hrs and 24 hrs post-injection. The 6 hrs and 24 hrs timepoints are measured using two separate cohorts of mice. For the 6 hrs timepoint: Saline, *n* = 8; 25 µg/kg LPS, *n* = 6; 50 µg/kg LPS, *n* = 7 mice. For the 24 hrs timepoint: Saline, *n* = 4; 25 µg/kg LPS, *n* = 4; 50 µg/kg LPS, *n* = 4 mice. Data is represented as mean ± SD. Differences between groups were compared using two-way ANOVA. **\*** ≤ 0.05, ** ≤ 0.01, *** ≤ 0.001, **** ≤ 0.0001.

Unexpectedly, LPS challenge did not alter frequencies of decidual Mo or Mo/Mφ cells expressing IFNγ (Figure 6A, 6B). Further, LPS did not alter IFNγ levels in Mo or Mo/Mφ (Figure 6C, 6D). However, cells residing within the CD3/CD19 dump-gate (admixture of T and B cells) did show increased MFI levels of IFNγ following LPS challenge at 6 and 24 hrs post treatment (Supplemental Figure 5A, 5B). Focusing on intracellular IFNγ in uNK subpopulations, LPS treatment did not affect trNK IFNγ^+^ frequencies nor IFNγ levels (Figure 6E-H). However in cNK, low dose LPS resulted in increased frequencies of IFNγ^+^ cells 24 hrs post challenge, whereas at 6 hrs both low and moderate doses of LPS led to modest increases in IFNγ levels, though these increases were not significant (Figure 6F-H). To summarize, LPS challenge consistently induced early production granzyme B in immature Mo and mature Mo/Mφ, as well as cNK. While LPS had antagonistic effects on TNFα production in Mo and Mo/Mφ, LPS potentiated TNFα expression in both trNK and cNK subtypes 24 hrs following challenge.

**Figure 6.**
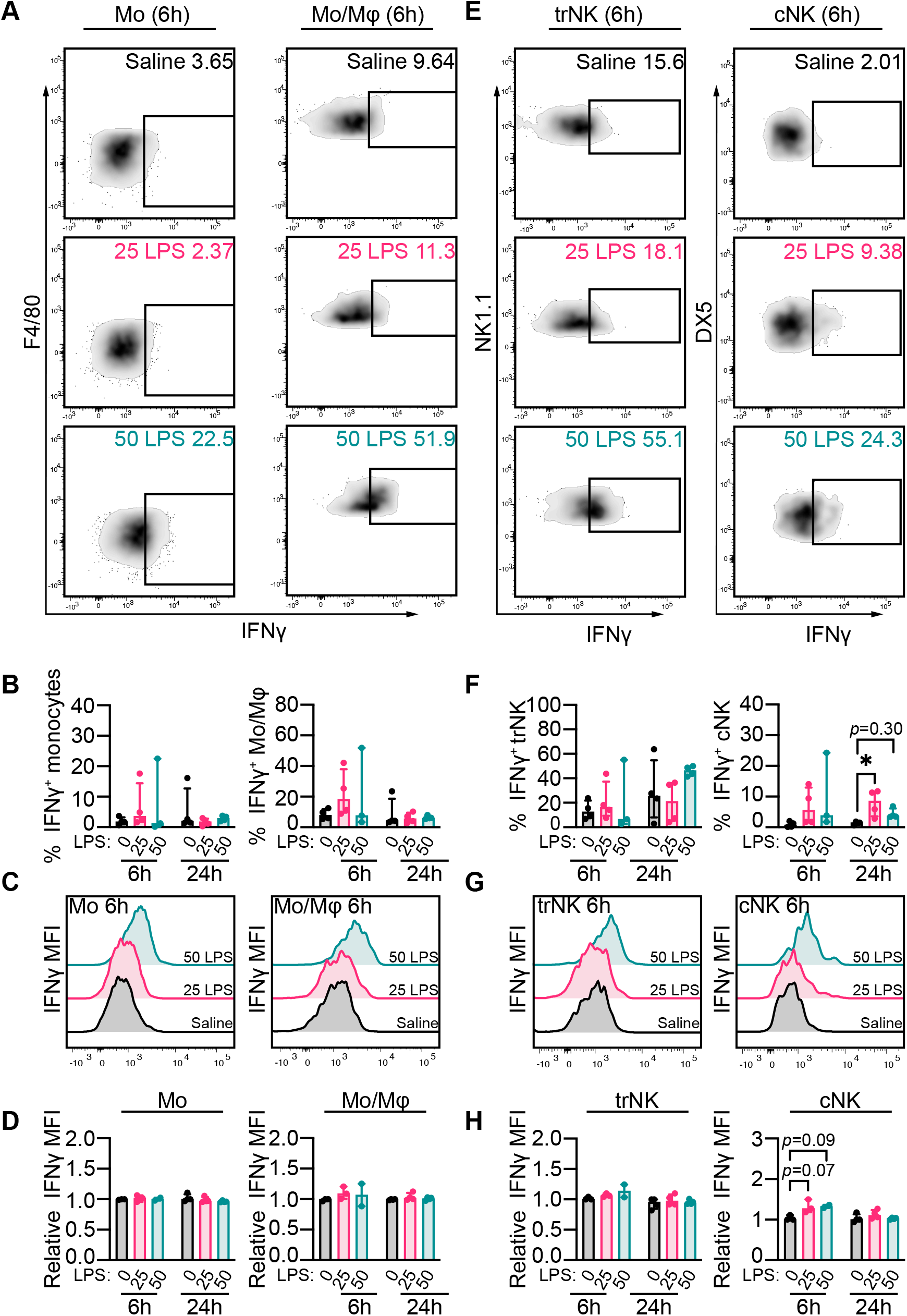
Temporal kinetics of uterine myeloid and NK intracellular IFNγ levels following LPS challenge. (A) Representative gating for expression of intracellular IFNγ 6 hrs following exposure to LPS (Saline, 25 µg/kg, 50 µg/kg) in immature monocytes (Mo), mature monocytes/Mφ (Mo/Mφ), trNK, and cNK in the decidua. (B) Representative trNK and cNK expression of IFNγ and TNFα, 6 hrs post-injection. Biological control for trNK and cNK expression of IFNγ and TNFα is non-cultured cNK. (C) Median fluorescent intensity (MFI) of IFNγ and TNFα in trNK and cNK (values relative to a Saline control for each experimental round), 6 hrs and 24 hrs post-injection. The 6 hrs and 24 hrs timepoints are measured using two separate cohorts of mice. For the 6 hrs timepoint: Saline, *n* = 8; 25 µg/kg LPS, *n* = 6; 50 µg/kg LPS, *n* = 7 mice. For the 24 hrs timepoint: Saline, *n* = 4; 25 µg/kg LPS, *n* = 4; 50 µg/kg LPS, *n* = 4 mice. Data is represented as mean ± SD. Differences between groups were compared using two-way ANOVA. **\*** ≤ 0.05, ** ≤ 0.01, *** ≤ 0.001, **** ≤ 0.0001.

## DISCUSSION

Inflammation stemming from infection or chronic inflammatory conditions in early pregnancy associate with pregnancy complications ranging from fetal growth restriction to fetal death. Yet it remains unclear the specific biological processes involved in inflammation induced poor pregnancy outcomes. In this study, we sought to understand how inflammation in early pregnancy impacts the composition or activity of key immune cells in the uterine environment. We show that LPS induces an expansion of cytolytically active immature monocytes that is followed by an expansion and activation of conventional NK that produce more cytolytic granules and pro-inflammatory cytokines. These LPS-induced alterations in uterine immune cell composition/activity associate with either enhanced decidual artery remodeling or fetal resorptions. This work provides new insight into the kinetics and activities of key myeloid and NK populations in the uterus following inflammatory insult in early pregnancy.

To model different severities of infection in early pregnancy, we injected pregnant mice with LPS, a well-established inflammatory stimulant. We confirmed there were increased levels of pro-inflammatory cytokines in the serum and increased pro-inflammatory gene expression in uterine tissue following LPS exposure. LPS has previously been shown to induce dose-dependent impairments in fetal outcome (i.e., preterm birth and neurological impairments) when administered near term in pregnant mice [59]. Indeed, low-dose LPS induced resorption in only 2 of 9 pregnancies, compared with 7 of 12 pregnancies with moderate-dose and 9 of 11 pregnancies with high-dose LPS. Though LPS increased readouts of peripheral and uterine inflammation, some of these increases were not dose-dependent. Other readouts of inflammation may better highlight a dose-dependent response to LPS. For example, LPS has previously been shown to increase *Il10* and *Tlr4* mRNA in hiPSC-CM cells in a dose-dependent manner [60]. Consistent with this, we show that LPS increases serum TNFα in a dose-dependent manner. Therefore, while administration of LPS in discreet concentrations produced expected rates of fetal resorption, the induction of the molecular machinery we examined does not tightly align with LPS dose suggesting that LPS-driven responses of many factors do not follow dose-dependent responses.

LPS-induced fetal resorption was bimodal in nature; uterine tissue on E9.5 was either fully dead or seemingly healthy. This bimodality of fetal demise following LPS challenge is consistent with the concept of quorum sensing and colonization resistance described in intestinal microbiota [61, 62]. Similarly, Muldoon et al. proposed a revised model of Mφ activation, in which LPS stimuli resulted in either low or high activation states in genetically identical Mφ culture; this bimodality of LPS-induced Mφ activation was largely attributed to ‘quorum licencing’, whereby cell density and external milieu determines a Mφ’s response to inflammatory stimulus [63, 64]. We examined gross fetal and placental metrics in the surviving pregnancies. Of those pregnancies that did not succumb to fetal resorption following low- and moderate-dose LPS exposure, LPS had no effect on fetal metrics at term and caused nuanced alterations in placental morphometry at term. It was unexpected to see no alterations in gross fetal or placental metrics despite clear induction of TNFα production in uterine immune cells. LPS exposure (and the resulting TNFα secretion) is widely known to induce preterm labour and fetal growth restriction [65–71]. Some differences between our study and other studies showing LPS-induced fetal/placental growth restrictions are the pregnancy model used (i.e., hetergeneic vs. monogeneic pregnancies), timing of LPS administration (embryonic day), LPS dosage, serotype of LPS administered, and mode of LPS administration (intraperitoneal vs. subcutaneous). We observed no suppression of placental growth at term, but this does not tell us if the placenta was functionally impaired. To estimate placental efficiency at term, we calculated birth weight to placental weight ratio (BW:PW) [72, 73]; we saw no differences in these measurements. However, a recent review by Christians, et al clarify that a BW:PW measurement does not always accurately indicate placental function/efficiency [74].

Inflammatory insult such as infection triggers a protective inflammatory response [75], and cytokine TNFα is a key regulator of inflammation, initiating several intracellular signalling cascades [76, 77]. Importantly, other models of LPS-induced fetal growth restriction and fetal death are dependent on TNFα [28, 58]. Indeed, LPS-challenge increased *Tnf* expression in uterine tissue, yet we observed fewer TNFα^+^ monocytes and Mφ 6 and 24 hrs following LPS injection, and these cells expressed lower levels of TNFα intracellularly. We propose two possible explanations for why we are observing decreased myeloid intracellular levels of TNFα following LPS-induced inflammatory challenge: First, LPS may have negative regulation of *Tnf* production [78]. LPS stimulus coordinates a complex intracellular response in Mφ: while there are positive feedback dominant loops in NF-κB expression [79], there are other pathways that constrain Mφ response to LPS (e.g., via reducing *Tnf* stability and translation) that help immune cells balance pathogen clearance and tissue damage [63,64,80,81]. Second, our observation that LPS-challenge increased *Tnf* mRNA expression in uterine tissue, yet decreased decidual trMφ expression of TNFα, suggests that decreased TNFα levels post-LPS challenge may be due to Mφ depletion of TNFα stores post-LPS stimulus. To address these possibilities, a dose response and time course would be needed to optimize our LPS stimulations and harvests.

Uterine innate immune activity is imperative for pregnancy success, participating in events from implantation, trophoblast invasion, artery remodeling, and decidualization [82–89]. Therefore, we assessed how LPS-induced inflammation affects key innate immune populations in the uterus. We observed a rapid expansion of immature monocytes in the decidual-fetal interface measured 6 hrs post-LPS challenge that likely reflected an influx of cytotoxic immature monocytes from the periphery to the uterus. A more comprehensive analysis of both peripheral and uterine myeloid populations would need to be performed to confirm the derivation of this expanding monocyte population. Interestingly, this rapid expansion of immature monocytes 6 hrs after LPS stimulus appeared to have been resolved by 24 hrs post-challenge. The rapid expansion of immature monocytes we observed is consistent with documentation that monocytes are one of the first-responding cells in inflamed tissue [90, 91]. Monocytes secreting chemokines that can alter the immune cell milieu at the decidual-fetal interface [92] and recruit other monocytes and NK from peripheral sources [67,72,93]. Indeed, we observed an expansion of cNK in the uterus in LPS-treated mice, and these cNK expressed more that expressed both granzyme B, TNFα, and IFNγ than control mice. This expansion of conventional NK to the decidual-fetal interface 24 hrs after LPS challenge, while non-significant, is nonetheless consistent with this mechanism for monocyte-mediated NK recruitment. Proportions of other immune populations (e.g., trNK, Mo/Mφ) decrease in response to LPS challenge, but a quantitative flow cytometry approach would be required to determine whether these populations are decreasing only relative to monocyte/cNK expansion or if their cell numbers are indeed decreasing in the uterus. Nonetheless, both trNK, Mo/Mφ were more cytolytically active 6 hrs following LPS challenge than those of control mice and these trNK expressed more TNFα at both timepoints. This early activation of trNK and Mo/Mφ indicate that these populations might have been present in the decidua prior to LPS exposure.

NK have been shown to be necessary for the pathogenesis of LPS-mediated fetal resorption [27,44,94]; however, the study showing requirement of NK in LPS-mediated fetal resorption did not examine the requirements of specific uNK subpopulations nor did they clarify the kinetics or activities of these populations. We show that LPS leads to heightened cNK expression of IFNγ, and heightened TNFα expression and cytotoxicity in both trNK and cNK. Conventional uNK secrete IFNγ and cytolytic granules to facilitate artery remodeling for a physiological pregnancy [56,82–85,87– 89,95,96]. Conversely, aberrant monocyte/NK secretion of cytotoxic granules (or secretion of these granules at the wrong developmental timepoint) may facilitate fetal resorption [52,97–100]. Also, uNK secretion of cytolytic granules has been shown to induce apoptosis of placental cells in human pregnancies [98, 101]. Indeed, the increase in immune cytotoxic activity in response to LPS challenge correlated with a temporal enhancement of decidual artery remodeling in some mice but correlated with increased fetal death in others. It remains unclear if the nature of response (i.e., enhanced arterial remodeling or increased resorption) to LPS challenge is predetermined or if an additional unknown factor(s) is/are involved in increasing susceptibility to LPS challenge. Of note, while we know uterine myeloid and NK are producing more cytolytic granules in response to our LPS-challenge, we do not know if these cells are depositing these granules; to test cytolytic deposition of these cells, a killing assay is required. Mo/Mφ are also known to mediate artery remodeling, mainly through phagocytosis of apoptotic smooth muscle cells [102]. However, we observed a heightened cytolytic activity post-LPS challenge in these myeloid populations. It remains unclear how this heightened Mo/Mφ cytotoxicity might affect Mo/Mφ functionality with regards to decidual artery remodeling, a macrophage-specific gene knockout experiment is needed to test this.

When performing flow cytometry experiments on mouse uterine immune cells, the limited available cell count and viability restrict the breadth of markers you can examine. We decided to focus this study on key myeloid and NK populations for pregnancy health and their roles in fetal resorption [26,52,84,88,97,98,100,104], however this does not rule out the importance of other immune populations. T cells, NK-T cells, innate lymphoid cells, and trophoblasts all have roles in pregnancy and may play important roles in LPS-mediated uterine inflammation [1–3,105–116]. Early stages of uterine blood vessel growth, branching, pruning, and remodeling are control. It will be important to study the effects of inflammation on these immune populations in the future. And while this study detangled some key effectors of inflammation-induced fetal resorption/artery remodeling, Knockouts/depletions of these key immune effectors (i.e., anti-asialo GM1 depletion of cNK [117]) and comprehensive examination of immune pathways involved (e.g., single cell sequencing and culture experiments) will be needed to fully understand the immune kinetics and activities driving inflammation-induced pregnancy outcomes.

Infections in pregnancy can be detrimental to the developing fetus, so this work set out to better understand the etiology of inflammation-induced fetal demise. We show that LPS challenge induces an expansion of immature monocytes in the uterus, followed by an expansion of cytolytic granule-producing cNK, likely contributing to LPS-mediated enhanced artery remodeling and/or fetal resorption. These findings help to clarify the immunological changes resulting from exposure to distinct grades of endotoxin challenge, providing insight into how inflammation shapes the decidual innate immune cell landscape.

## ACKNOWLEDGEMENTS

We would like to extend our gratitude to Dr. Julian Christians, Simon Fraser University, for providing us with LPS. We would like to thank Dr. Louis Lefebvre, University of British Columbia, for his helpful discussions in the design of the DBA/C57BL/6 breeding strategy. We would also like to thank Dr. Joannie Allaire and Dr. Bruce Vallance, University of British Columbia, for providing RNA from inflamed mouse gut tissue and for helping us to optimize histological stains. We would like to thank Dr. Laura Sly, University of British Columbia, for her helpful discussions central to data interpretation. Lastly, we would like to thank members of the Beristain Lab; Matthew J Shannon, Jasmin Wächter, and Burak Koksal, for their aid in editing this manuscript. Graphical abstract created with BioRender.com.

## AUTHOR CONTRIBUTIONS

AGB, LES, and BC designed the research. LES, BC, and JB performed the experiments. LES, BC, and AGB analysed the data and wrote the paper. All authors read and approved the manuscript.

---

**Supplemental Figure 1.**
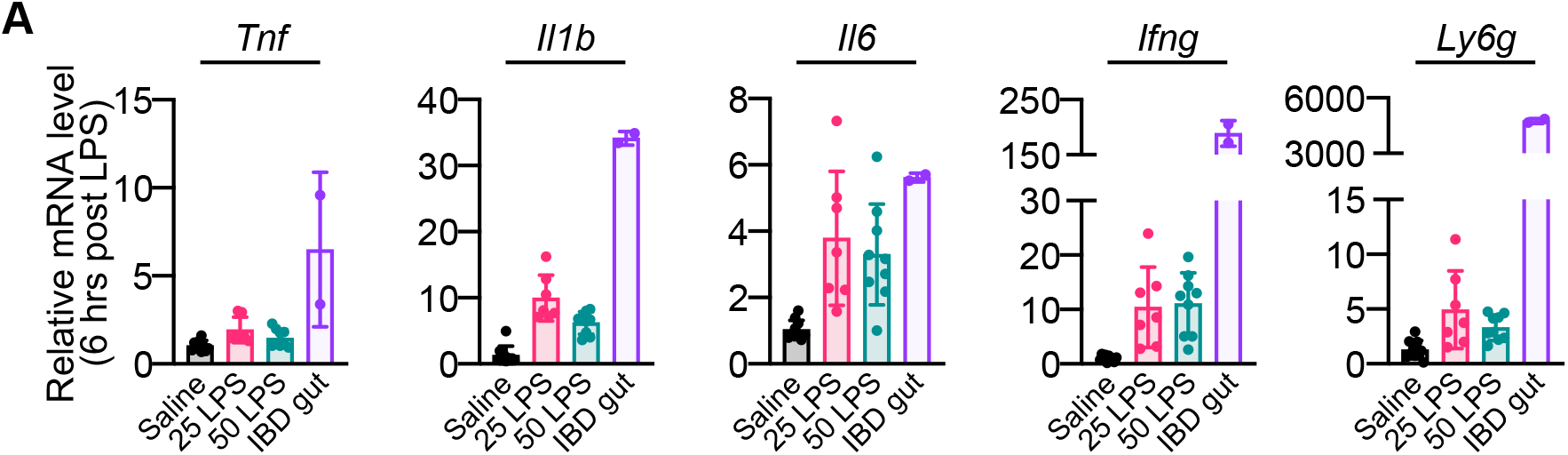
qPCR with inflamed mouse gut positive control. (A) Gene expression levels of *Tnf*, *Il1b*, *Il6*, *Ifng*, *Ly6g* within uterine tissue 6 hrs after injection. Saline, *n* = 8; 25 µg/kg LPS, *n* = 6; 50 µg/kg LPS, *n* = 7. control. Data is represented as mean ± SD; no statistical analyses were performed.

**Supplemental Figure 2.**
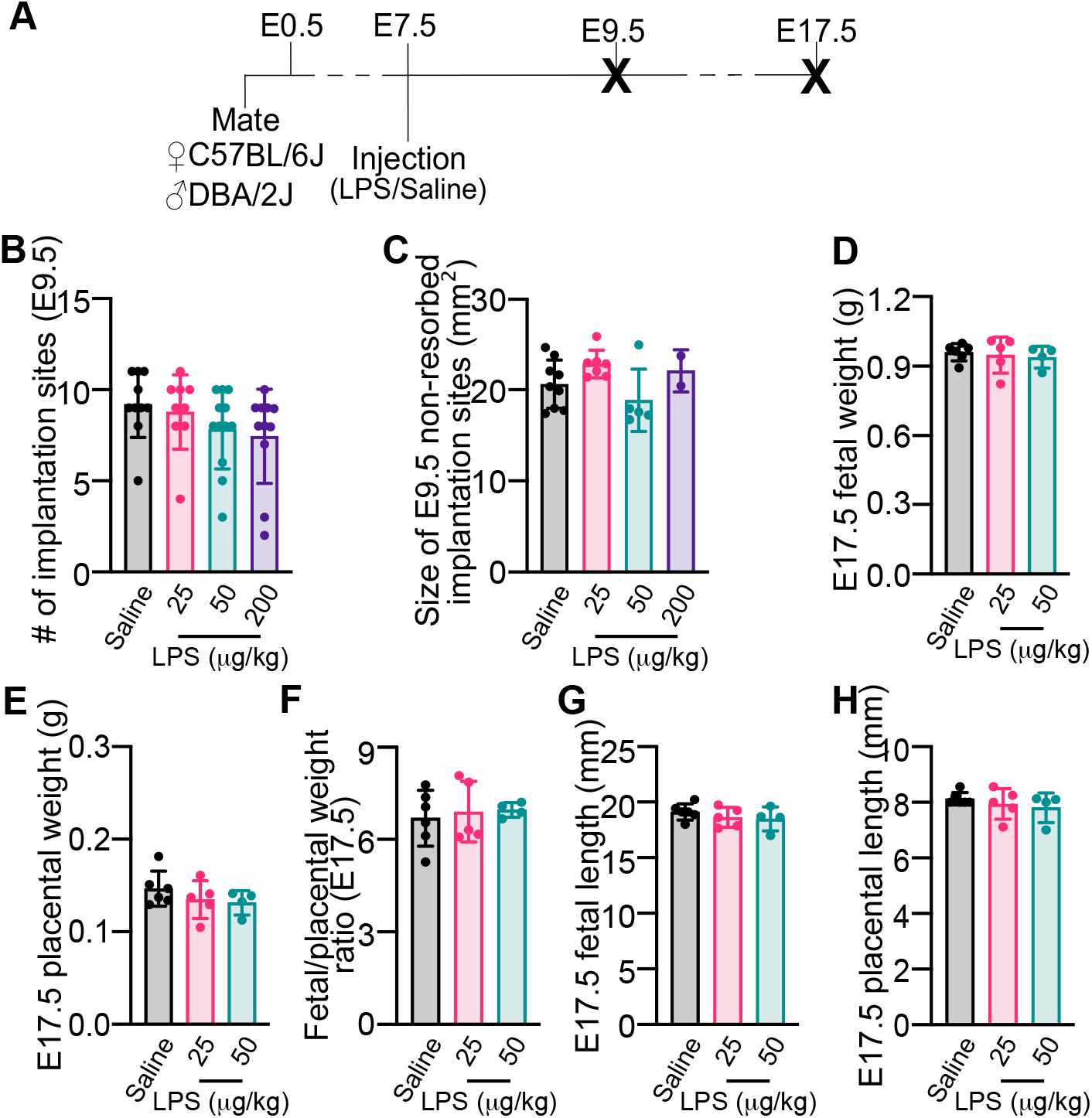
LPS does not affect term fetal/placental metrics. (A) Schematic of timeline for mouse mating, injection, and sacrifice (bold **X**) for Figure 2 experiments. (B) Number of implantation sites per mouse measured 48 hrs following injection, and (C) area of live implantation sites per mouse 48 hrs following injection. Saline, *n* = 9; 25 µg/kg LPS, *n* = 7; 50 µg/kg LPS, *n* = 5; 200 µg/kg LPS, *n* = 2 mice. (D) Fetal weights, (E) placental weights, (F) fetal: placental weight ratios, (G) fetal crown-rump lengths, and (H) placental diameter of non-resorbed implantation sites at E17.5. Saline, *n* = 6; 25 µg/kg LPS, *n* = 5; 50 µg/kg LPS, *n* = 4 mice. Data is represented as mean ± SD. Differences between groups were compared using one-way ANOVA. **\*** ≤ 0.05, ** ≤ 0.01, *** ≤ 0.001, **** ≤ 0.0001.

**Supplemental Figure 3.**
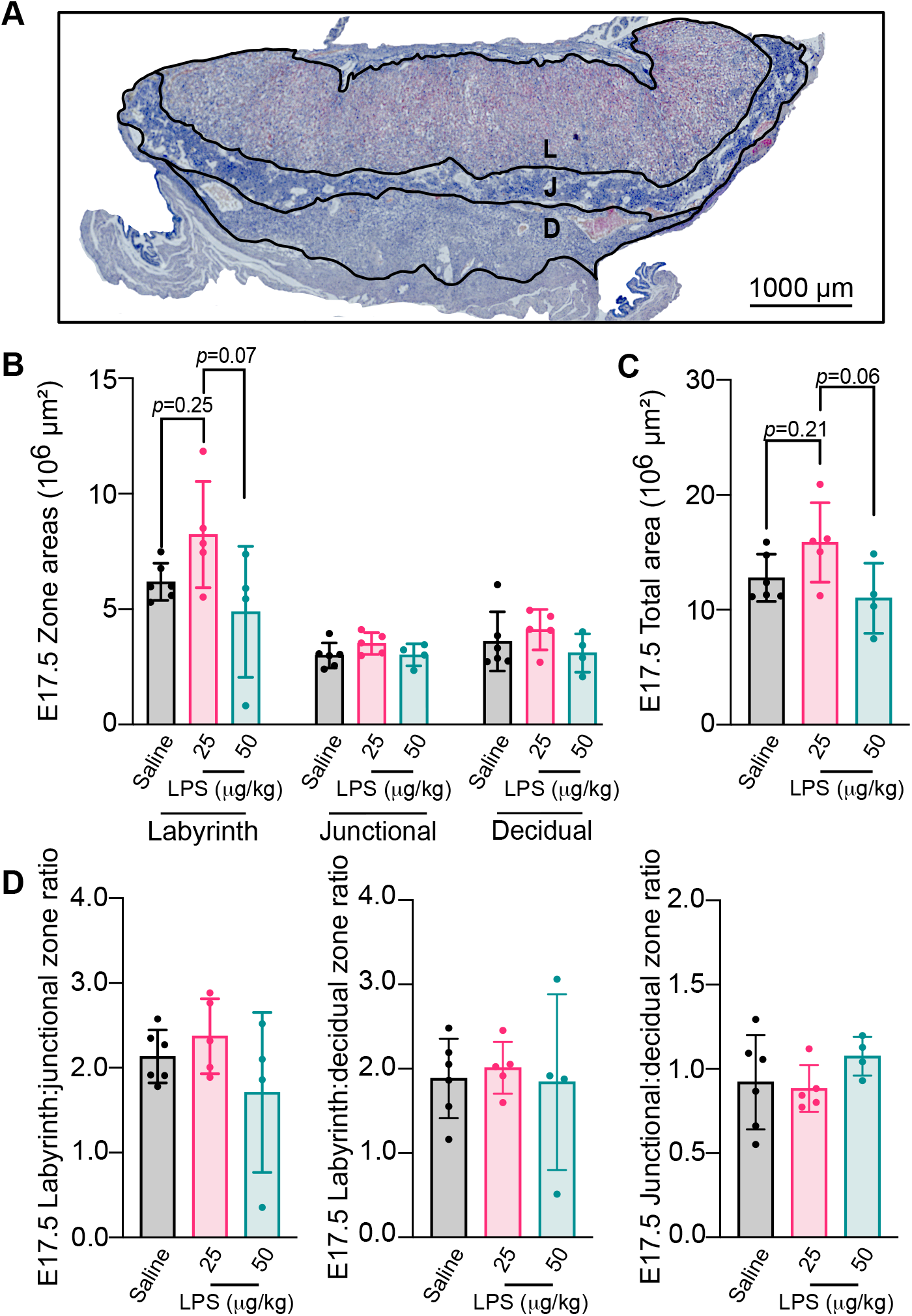
LPS does not affect placental morphometry of surviving pregnancies. (A) Representative image of cross section of implantation site at E17.5, and perimeters of placental zones (labyrinth, L; junctional, J; decidual, D). Scale bar = 1000 µm. (B) Areas of placental zones (labyrinth, junctional, decidual), (C) total implantation site area (sum of labyrinth, junctional, and decidual zones), and (D) placental zone ratios (labyrinth: junctional, labyrinth: decidual, junctional: decidual) all at E17.5. Saline, *n* = 6; 25 µg/kg LPS, *n* = 5; 50 µg/kg LPS, *n* = 4 mice. Data is represented as mean ± SD. Differences between groups were compared using one-way ANOVA. **\*** ≤ 0.05, ** ≤ 0.01, *** ≤ 0.001, **** ≤ 0.0001.

**Supplemental Figure 4.**
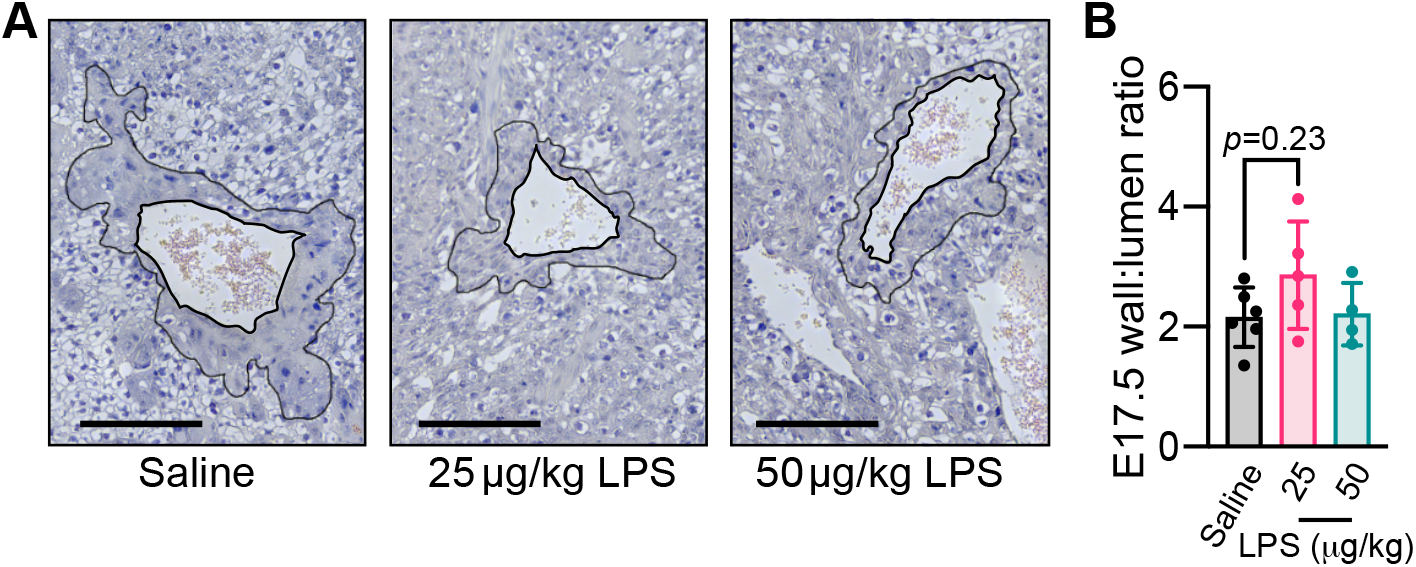
LPS does not affect artery remodeling in late pregnancy. (A) Representative images of E17.5 arteries stained for hematoxylin and eosin. Scale bar = 100 µm. (B) Artery wall:lumen ratio per mouse at E17.5. Saline, *n* = 6; 25 µg/kg LPS, *n* = 5; 50 µg/kg LPS, *n* = 4 mice. Data is represented as mean ± SD. Differences between groups were compared using one-way ANOVA. **\*** ≤ 0.05, ** ≤ 0.01, *** ≤ 0.001, **** ≤ 0.0001.

**Supplemental Figure 5.**
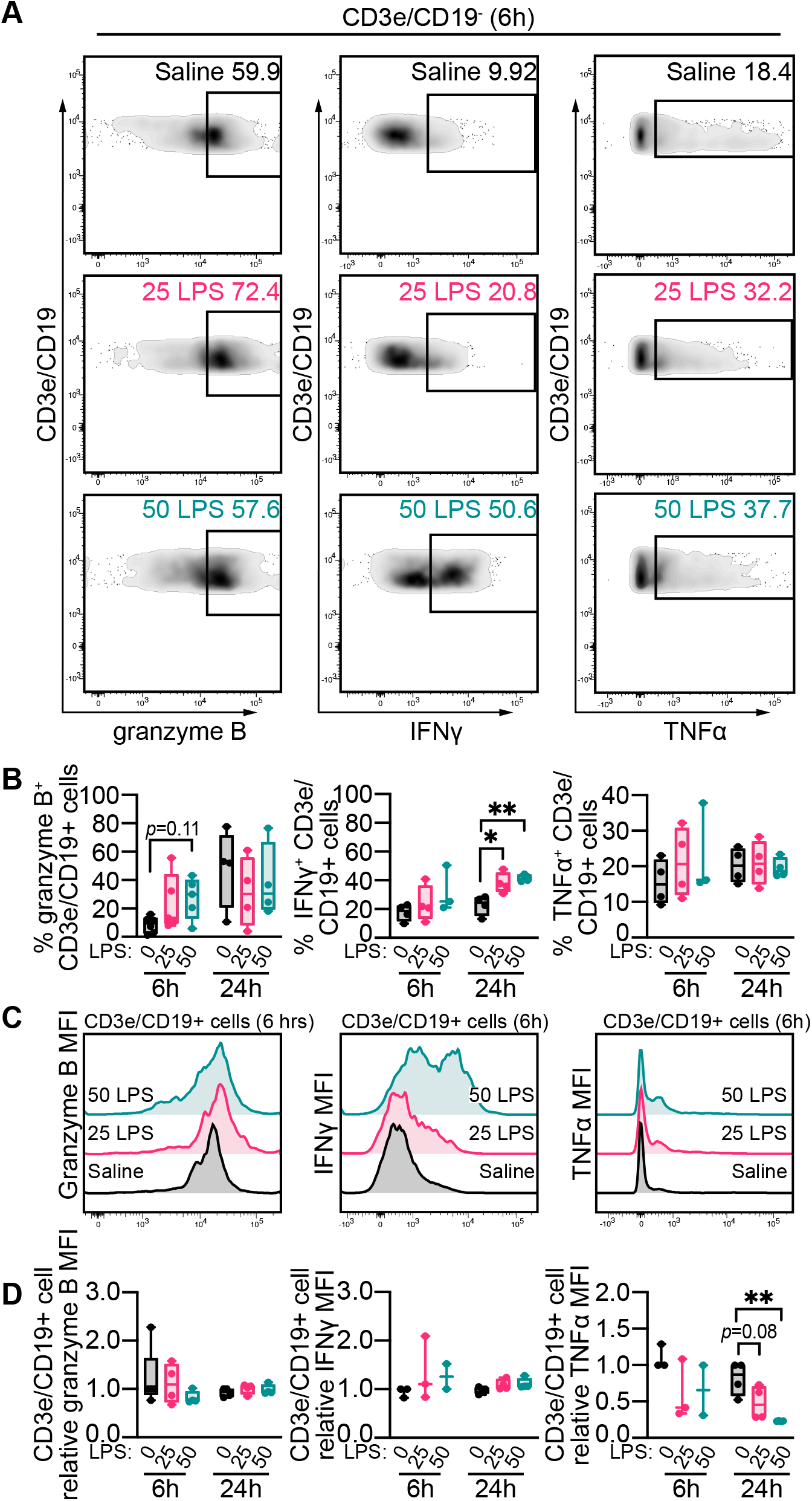
Temporal kinetics of uterine T/B intracellular cytolytic and pro-inflammatory factor levels following LPS challenge. (A) Representative gating for T/B cell expression of intracellular granzyme B, IFNγ and TNFα 6 hrs following exposure to LPS (Saline, 25 µg/kg, 50 µg/kg). (B) Proportions of T/B cells expressing granzyme B, IFNγ and TNFα 6 hrs and 24 hrs post-injection. (C) Representative T/B cell expression of granzyme B, IFNγ and TNFα, 6 hrs post-injection. Pos. control for granzyme B expression is stimulated T/B cells. Neg. control for IFNγ and TNFα expression is non-cultured T/B cells. (D) Median fluorescent intensity (MFI) of granzyme B, IFNγ and TNFα in T/B cells (values relative to a Saline control for each experimental round), 6 hrs and 24 hrs post-injection. The 6 hrs and 24 hrs timepoints are measured using two separate cohorts of mice. For the 6 hrs timepoint: Saline, *n* = 8; 25 µg/kg LPS, *n* = 6; 50 µg/kg LPS, *n* = 7 mice. For the 24 hrs timepoint: Saline, *n* = 4; 25 µg/kg LPS, *n* = 4; 50 µg/kg LPS, *n* = 4 mice. Data is represented as mean ± SD. Differences between groups were compared using two-way ANOVA. **\*** ≤ 0.05, ** ≤ 0.01, *** ≤ 0.001, **** ≤ 0.0001.

**Supplemental Table 1. Raw mesoscale data shown in Figure 1B.**

**Supplemental Table 2. Raw flow cytometry data including mouse excluded from study.**

